# A Novel *in vivo* Model of Anaerobic Infection: The Investigation of *Clostridium perfringens* in *Galleria mellonella* Larvae

**DOI:** 10.1101/382077

**Authors:** Sammy Kay, Joseph Edwards, Joseph Brown, Ronald Dixon

## Abstract

**ABSTRACT:** Important research progress into the mechanisms of *Clostridium perfringens* associated diseases (CPAD) has been slowed by the lack of a reliable infection model. Wax moth larvae (*Galleria mellonella*) have emerged as a viable alternative to traditional mammalian organisms since they are economic, survive at 37°C and require no specialist equipment. This study aims to establish whether *G. mellonella* larvae can be developed as a viable model for the study of CPAD and their suitability for studying novel treatment strategies. In addition, the study demonstrates a novel time-lapse approach to data collection. Mortality and morbidity rates of larvae challenged with 10^5^ CFU of *C. perfringens* isolates from various sources were observed over 72h and dose response data obtained using inoculum sizes of 10 - 10^5^ CFU. Phenoloxidase enzyme activity was investigated as a marker for immune response and tissue burden by histopathological techniques. Results show that *C. perfringens* is pathogenic towards *G. mellonella* although potency varies between isolates. Infection activates the melanisation pathway resulting in melanin deposition but no increase in enzyme activity was observed. Efficacy of antibiotic therapy (penicillin G, bacitracin, neomycin and tetracycline) administered parenterally loosely correlates with that of *in vitro* analysis. The findings suggest *G. mellonella* can be a useful *in vivo* model of infection when investigating CPAD. Although they are unlikely to replace traditional mammals they may be useful as a pre-screening assay for virulence of *C.perfringens* strains or as a simple, cheap and rapid *in vivo* assay in the development and pre-clinical development of novel therapeutics.

**Highlights:**

- Novel *in vivo* model for the study of *Clostridium perfringens* infection.
- Novel time-lapse approach to data collection.
- First report of the use of *G. mellonella* model for characterizing virulence in *C. perfringens* strains.
- Antibiotic therapy in the model that loosely correlates with *in vitro* testing.

## Introduction

Insect models have been shown to be helpful in our understanding of the virulence of bacterial pathogens in humans (Tsai *et al.*, 2016). Insects share some similarities with mammalian processes and possess a basic innate immune system (Ramarao *et al.*, 2012a). Non-vertebrates are not associated with the same ethical considerations as the use of mice or other vertebrates (Peterson *et al.*, 2008). Although some alternative invertebrate models of infection have been developed i.e. *Bombyx mori* (silkworm), *Manduca sexta* (tobacco hornworm), *Drosophila melanogaster* (fruit fly), *Danio renio* (zebra fish) and *Caenorhabditis elegans* (nematode) (Browne *et al.*, 2013); the larval stage of the Greater Wax moth (*Galleria mellonella*) has emerged as an insect model of particular value since it survives at 37°C, which is essential for demonstration of specific microbial virulence factors (Smoot *et al.*, 2001).

The larvae are inexpensive to obtain and easy to maintain using basic equipment (Ramarao *et al.*, 2012a). The model does not require ethical approval and coupled with fast reproduction time, allows for a high-throughput of experiments compared with mammal systems. Whilst *G. mellonella* has not evolved an adaptive immune response, they possess a semi-complex cellular and humoral innate immunity. This innate system, in insects, shares remarkable similarities to that of mammals (Jander *et al.*, 2000). The cellular immune response in the larvae consists of haemocytes which play a role in phagocytosis, encapsulation and clotting as an antimicrobial response (Tojo *et al.*, 2000). The humoral response consists of various antimicrobial peptides, opsonins, extracellular nucleic acid traps and the phenoloxidase pathway as described in detail recently by Tsai *et al.*, (2016).

Development of infection models with *G. mellonella* larval hosts has involved a diverse range of microbes including: *Cryptococcus neoformans* (Mylonakis *et al.*, 2005), *Burkholderia cepacia* (Seed and Dennis, 2008), *Yersinia pseudotuberculosis* (Champion *et al.*, 2009), *Acinetobacter baumannii* (Peleg *et al.*, 2009), *Campylobacter jejuni* (Senior *et al.*, 2011), *Candida albicans* (Brennan *et al.*, 2002), *Legionella pneumophilia* (Harding *et al.*, 2013), *Pseudomonas aeruginosa* (Beeton *et al.*, 2015), *Mycobacterium fortucinium* (Entwistle and Coote, 2018) and *Vibrio parahaemolyticus* (Wagley *et al.*, 2018).

In the present study, we have investigated the use of *G. mellonella* larvae as a potential *in vivo* infection model for the study of complex and multifactorial CPAD. *C. perfringens* is a ubiquitous Gram-positive, spore-forming, anaerobic bacilli, classically characterised by four major extracellular toxins: alpha (*cpa*), beta (*cpb*), epsilon (*etx*) and iota (iA). Recently necrotic enteritis like B toxin (*netB*) and enterotoxin (*cpe*) has been added to the classification system (Table 1)(Rood *et al.*, 2018). The bacteria produce a plethora of non-typing toxins including haemolysin, perfringolysin O (PFO), collagenase (*colA*) (Songer, 1996), beta2 toxin (*cpb2*) (Van Asten *et al.*, 2010), and binary enterotoxin A and B (*becA and becB*) (Yonogi *et al.*, 2014) with differing and elusive roles in the bacterial mode of infectivity.

**Table 1:**
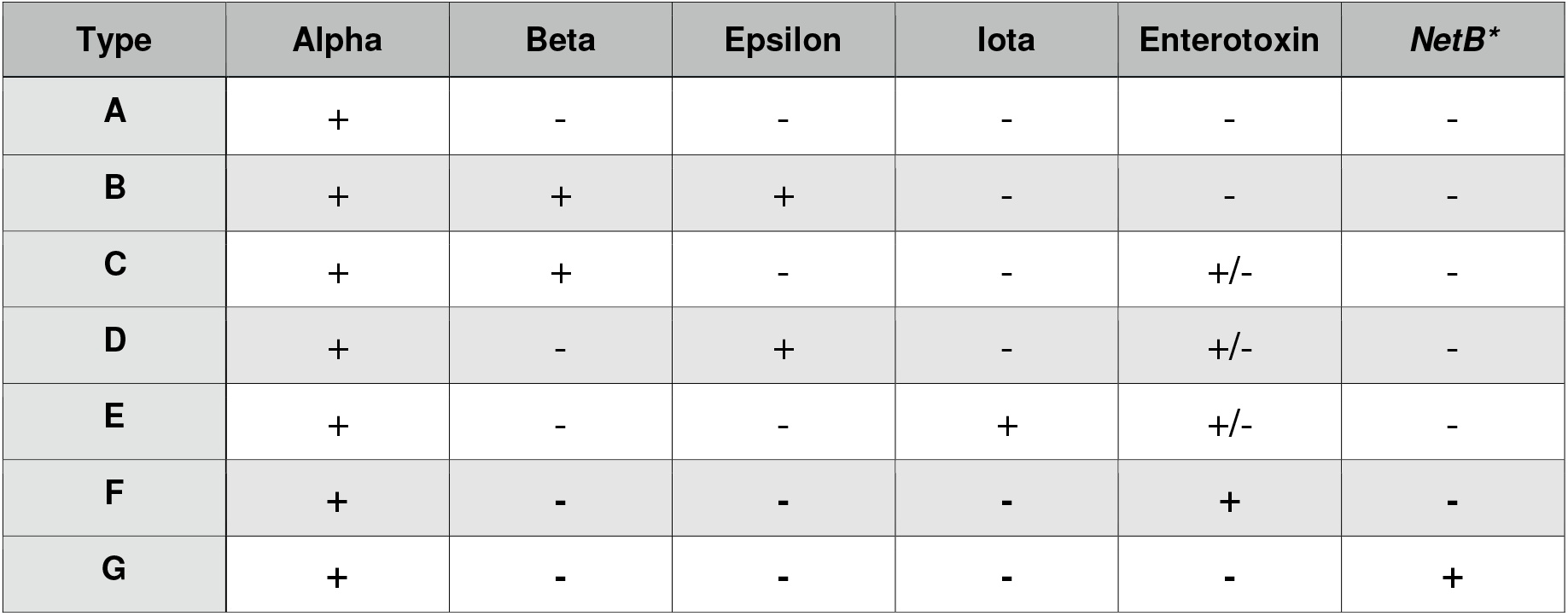
Toxin type classification system for *C. perfringens*. Isolates are characterised by the production of one, or more, of the now six major toxins: alpha, beta, epsilon, iota, *necrotic enteritis like B and enterotoxin. Alpha toxin is produced by all types of *C. perfringens*. (Rood *et al.*, 2018)

| Type | Alpha | Beta | Epsilon | Iota | Enterotoxin | *NetB* * |
| --- | --- | --- | --- | --- | --- | --- |
| A | + | - | - | - | - | - |
| B | + | + | + | - | - | - |
| C | + | + | - | - | +/- | - |
| D | + | - | + | - | +/- | - |
| E | + | - | - | + | +/- | - |
| F | + | - | - | - | + | - |
| G | + | - | - | - | - | + |

*C. perfringens* is one of the leading causes of food poisoning in the UK and worldwide. It is implicated in 80-95% of reported gas gangrene cases (Titball *et al.*, 2005) and causes large economic losses in the agricultural industry (Wade and Keyburn, 2015). Research into this important bacterial pathogen has been slowed by the lack of standardised disease models for assessing virulence and *in vivo* susceptibility to antimicrobial drugs. The *G. mellonella* larvae model of infection offers a rapid insight into the virulence of pathogen isolates and an alternative economical screening method for progressing novel antimicrobial compounds prior to mammalian studies with the anaerobe.

To date, no detailed reports of *C. perfringens* infection models with *G. mellonella* have been published. It should be noted, that an unpublished report suggests alpha and enterotoxin from *C. perfringens* lack toxicity against the larvae. This present study investigates the extent of *G. mellonella* susceptibility to 19 distinct *C. perfringens* isolates including a standard reference strain and explores the response of the larvae to antibiotic therapy. In addition, we introduce a novel imaging capture and time-lapse methodology designed to increase accuracy whilst easily monitoring disease progression.

## Materials and Methods

### Bacterial strains and preparation of inoculum

*C. perfringens* was sourced from our culture collection and isolates selected based on toxin profiles (Table 2). Neonatal isolates were acquired from study at Imperial College London (Sim *et al.*, 2015). Cultures were stored in 30% glycerol (Fisher Scientific, UK), 70% Brain heart infusion stocking media (Oxoid, UK) at −80°C and cultured on BHI agar (Oxoid, UK). Liquid cultures were grown to mid-log phase, in BHI or thioglycollate broth (Oxoid, UK) and incubated at 37°C for hours. Bacterial inoculum were adjusted by absorbance with OD_620_ v CFU/mL calibration curve (data not shown). Cells were pelleted by centrifuging at 3170xg at room temperature for 2 minutes (Heraeus Megafuge 8) and washed once with 0.1% peptone water (Oxoid, UK). Dilutions were drawn into a 1mL syringe fitted with a 30g sterile needle (BD plastipak).

**Table 2:** *Clostridium perfringens* toxin profiles of strains and isolates used within this study.

| Isolate | Source | *cpa* | *cpb* | *etx* | *iA* | *cpe* | *netb* | *cpb2* | *pfoA* | *tpeL* | *colA* | *becA* | *becB* | *cna* | Type |
| --- | --- | --- | --- | --- | --- | --- | --- | --- | --- | --- | --- | --- | --- | --- | --- |
| Q143.CN7 | Neonatal | + | - | - | - | - | - | + | + | - | + | - | - | - | A |
| Q100.CN11 | Neonatal | + | - | - | - | - | - | - | - | - | - | - | - | - | A |
| Q215.CN2 | Neonatal | + | - | - | - | - | - | + | + | - | - | - | - | - | A |
| Q020.CN1 | Neonatal | + | - | - | - | - | - | - | - | - | + | - | - | + | A |
| JBFR014 | Free Range | + | - | - | - | - | - | - | - | - | - | - | - | - | A |
| ATCC 13124 | Type Strain | + | - | - | - | - | - | - | + | - | + | - | - | - | A |
| LPCP001 | Caprine | + | - | - | - | - | - | - | + | - | + | - | - | - | A |
| LPCP023 | Caprine | + | - | - | - | - | - | - | + | - | + | - | - | - | A |
| LPCP042 | Caprine | + | - | - | - | - | - | - | + | - | + | - | - | - | A |
| EDCB283 | Retail Chicken | + | - | - | - | - | + | + | + | - | + | - | - | - | G |
| EDCB093 | Retail Chicken | + | - | + | - | - | - | - | + | + | + | - | - | - | B |
| JBRC013 | Reptile | + | - | - | - | - | + | - | - | - | + | + | - | - | A |
| JBCNJ58 | Broiler | + | - | - | - | - | + | - | + | - | + | - | - | - | G |
| JBCNCR03 | Broiler | + | - | - | - | - | + | - | + | + | + | - | - | - | G |
| JBCNC055 | Broiler | + | - | - | - | - | - | + | + | - | + | - | - | - | A |
| JBCND021 | Broiler | + | - | - | - | - | + | + | + | - | + | - | - | - | G |
| JBCNI050 | Broiler | + | - | - | - | - | - | - | - | - | - | - | - | - | A |
| JBCND053 | Broiler | + | - | - | - | - | - | - | + | - | + | - | - | - | A |
| JBCNJ055 | Broiler | + | - | - | - | - | - | + | + | - | + | - | - | - | A |

### *Galleria mellonella* challenge assay

Larvae were purchased from Live Foods UK (Rooks Bridge, Somerset UK). Upon arrival, they were stored in wood shavings, at room temperature, in the dark. To avoid sampling biases, larvae with any signs of melanisation or deformity were rejected. Larvae were weighed and only larvae meeting the criteria of 250mg ± 50mg were utilised. Prior to all injections, all larvae were immobilized by submerging the 24 micro-well plate (Fisher Scientific, UK) in ice for 15 minutes. An automated syringe pump (KD Scientific, US) was supplied with the drawn 1mL syringes and set to administer 1μL/s for 10 seconds. A safety restraint device was used, outlined by Dalton *et al.*, (2017). To minimise adverse effects, larvae were injected into the right posterior proleg. A melanisation scoring system adapted from Senior *et al.*, (2011) is shown in Figure 1 and employed throughout. Results were obtained longitudinally by the use of a novel image capture system. Briefly, the system was constructed with a benchtop incubator, with transparent sides and top (Stuart Scientific SI140). Images were recorded by a Logitech 15MP camera coupled with time-lapse software (SkystudioPro). For each experiment, images of the larvae were recorded every 10 minutes for 72 hours.

**Figure 1:**
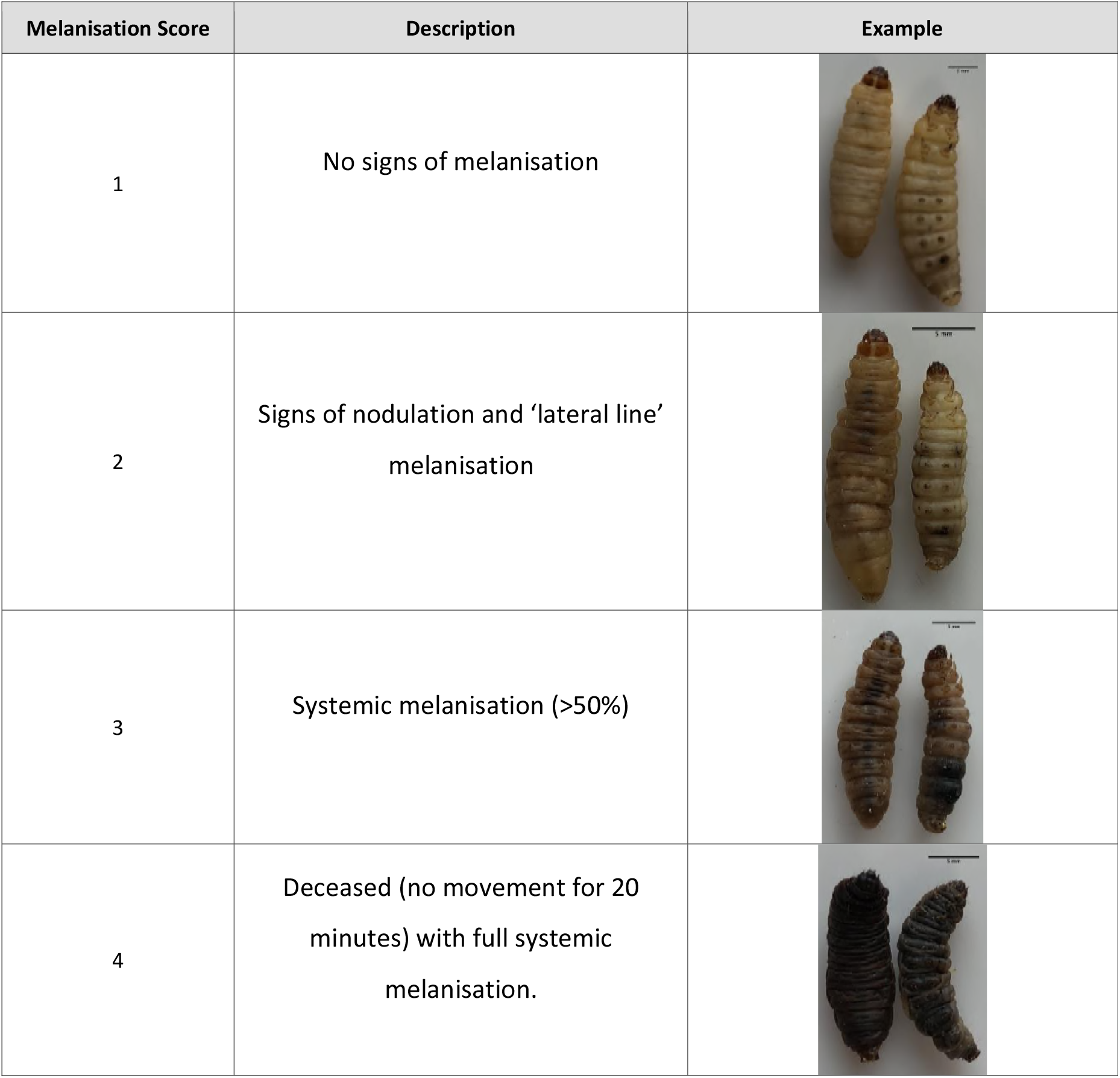
Melanisation scoring system adapted from Senior *et al.*, (2011).

### Pathogenicity screening

Nineteen *C. perfringens* environmental isolates from a variety of sources (Table 2) were prepared for inoculation as shown above. Ten larvae were challenged with 10μl of 10^7^ CFU/mL. Each experiment included ten larvae injected with 0.1% peptone water as controls. Larvae were placed in Petri dishes lined with greaseproof paper to allow for greater contrast in images. All groups were incubated aerobically at 37°C and survival recorded after 72 hours. The experiments were repeated in triplicate and an average survival result recorded.

### Dose dependent challenge

In dose dependent studies, larvae were prepared by placing one per well in 24 micro-well plastic plates (Fisher Scientific, UK). All larvae were treated identically to ensure injection and isolate continuity throughout. ATCC^®^ 13124^™^, JBFR014 and JBCNJ055 were investigated by preparing a dilution series of each isolate and larvae were injected with a dilution of 10μL of washed cultures ranging from approximately 10^3^ to 10^7^ CFU/mL. In addition, 24 larvae were injected with 10μL of 0.1% peptone water as negative controls. All six plates for each isolate were recorded simultaneously within the image recording system described previously. The larvae were incubated for 72 hours at 37°C. Images were taken every 10 minutes for 72 hours. Melanisation scores and mortality were recorded at 0,12, 24, 36, 48, 60 and 72 hours post injection. All experiments were repeated in triplicate and average scores recorded.

### Haemolymph extraction

Twelve larvae were challenged with 10μL of approximately 10^7^ CFU/mL of JBCNJ055 washed culture and twelve larvae injected with 10μL of 0.1% peptone water as controls. All larvae were incubated at 37°C simultaneously for 72 hours. At 24 hour intervals post challenge, three larvae from each group were placed into separate sterile 15mL centrifuge tubes (Fisher Scientific, UK) and submerged in ice for 15 minutes for immobilization. Haemolymph was extracted by removing the posterior two segments and bleeding into a sterile pre-chilled 1.5mL micro centrifuge tube (Eppendorf, UK). 25μL of extracted haemolymph was transferred into 75μL of 50mM PBS (pH 6.5) (Gibco, UK) then briefly vortexed and centrifuged to pellet the cells at 11,180xg for 10 minutes at 4°C. The supernatant was transferred into sterile 1.5mL tubes. Samples were processed within 15 minutes of extraction to avoid melanisation.

### Phenoloxidase activity

Phenoloxidase activity was measured using a microplate enzyme assay (Eleftherianos *et al.*, 2008). Briefly, a reaction mixture containing 115μL of 50 mM PBS (pH 6.5) and 10μL haemolymph plasma was prepared. 25μL of 20 mM 4-methyl catechol (Sigma, UK) was added as enzyme substrate and 2μL of 10mM *Escherichia coli* LPS (Sigma, UK) was added to controls. Plates were subjected to 1 hour of low agitation (25RPM) at room temperature to activate endogenous pro-phenoloxidase prior to addition of the substrate. The change in absorbance was read at 490nm for 1 hour at room temperature with a reading taken every 60 seconds using a plate reader (Fluostar Optima). Each reaction was repeated in triplicate.

### Histopathology of Galleria samples

Groups of three larvae were injected with 10μL of approximately 10^7^ CFU/mL of JBCNJ055 and incubated at 37°C for 72 hours. Control groups were injected with 10μL of 0. 1% peptone water. At 36h post-injection larvae were immobilised by submerging in ice for 15 minutes. Larvae were injected with 150μL of 10% neutral buffered formalin (Fisher Scientific, UK) (for internal fixation) until turgid and stored in 10% neutral buffered formalin for 36h prior to routine tissue processing. Larvae were laterally dissected and both halves embedded into wax blocks. Tissue sections (5μM) were cut by routine methods, mounted on glass slides and stained with either H+E, Masson Fontana, or Gram stains. Sections were imaged using light microscope (Zeiss Primostar) and a 1080p camera (Mitotic).

### MIC Determination

*C. perfringens* isolate JBCNJ055 was sub-cultured in 10mL of thioglycollate broth (Oxoid, UK) and incubated for 6 hours, aerobically at 37°C. Broth dilution MIC assays were produced in microtiter plates and prepared with ranges from 128μg/mL to 0.25μg/mL of penicillin G, bacitracin, tetracycline, gentamicin and neomycin. MICs were recorded as the lowest concentration of antimicrobial agent that completely inhibits growth turbidity compared with positive controls where antibiotics were omitted. MIC testing was repeated in triplicate on three separate occasions.

### Antibiotic Therapy

As positive controls, 24 larvae were challenged with 10μL of approximately 10^7^ JBCNJ055 culture followed by an injection of 10μL of sterile dH_2_O. A further 24 larvae were injected with 10μL of the same inoculum with a second 10μL injection of either penicillin G (2mg/kg), bacitracin (64mg/kg), tetracycline (64mg/kg), or neomycin (2400mg/kg). Injected concentrations are shown in Table 3. Treatment injections were administered within 15 minutes of the bacterial injection. Dosages were chosen to equate with greater than 10x *in vitro* MIC. All larvae were subjected to toxicity testing prior to therapy trials (data not shown). Finally, 24 larvae were injected with 10μL 0.1% peptone water as negative controls (Oxoid, UK). Larvae were incubated simultaneously at 37°C for 72 hours. Morbidity and mortality scores were recorded at 0, 12, 24, 36, 48, 60 and 72 hours post injection. All experiments were repeated in triplicate and average scores recorded.

**Table 3:** *In vitro* MIC results with associated therapeutic dosage. Survival percentages ± Standard deviation after 72 hours of being treated with 10μL injected doses of 10x *in vitro* MIC against 10μL injection of 10^7^ JBCNJ055. ‘+’ indicates no treatment administered. ‘-’ indicates infection with 0.1% peptone water as trauma controls.

| Treatment | *In vitro* MIC<br>(μg/mL) | > x10<br>(μg/mL) | Antibiotic<br>dosage x100<br>(mg/mL) | Larval<br>Dosage<br>(μg/larvae) | Larval Survival<br>(% ± SD) |
| --- | --- | --- | --- | --- | --- |
| + Control | | | | 0 | 73.61 ± 6.36 |
| - Control | | | | 0 | 100 ± 0 |
| Penicillin G | 0.18 | 5 | 5 | 5 | 68.06 ± 15.77 |
| Bacitracin | 1.56 | 15.6 | 1.6 | 16 | 94.45 ± 4.8 |
| Neomycin | 60 | 600 | 60 | 600 | 65.28 ± 18.78 |
| Tetracycline | 1.56 | 15.6 | 1.6 | 16 | 79.17 ± 14.43 |

### Statistical analysis

The data from each repeat was pooled. Analysis for dose dependent challenges was conducted using Kaplan-Meier survival distributions and tested for statistical significance using the Log rank (Mantel Cox) test (pooled over strata). Enzyme activity was analysed using Mann-Whitney U tests due to non-normal distribution in the data. All tests were carried out in SPSS, 24 (IBM).

## Results

### Isolate pathogenicity

Figure 2 highlights the variability seen in pathogenicity against *G. mellonella* with different isolates of *C. perfringens*. Isolates derived from human sources generally show lower pathogenicity than those of animal origin. There is no obvious correlation between toxin type and pathogenicity.

**Figure 2:**
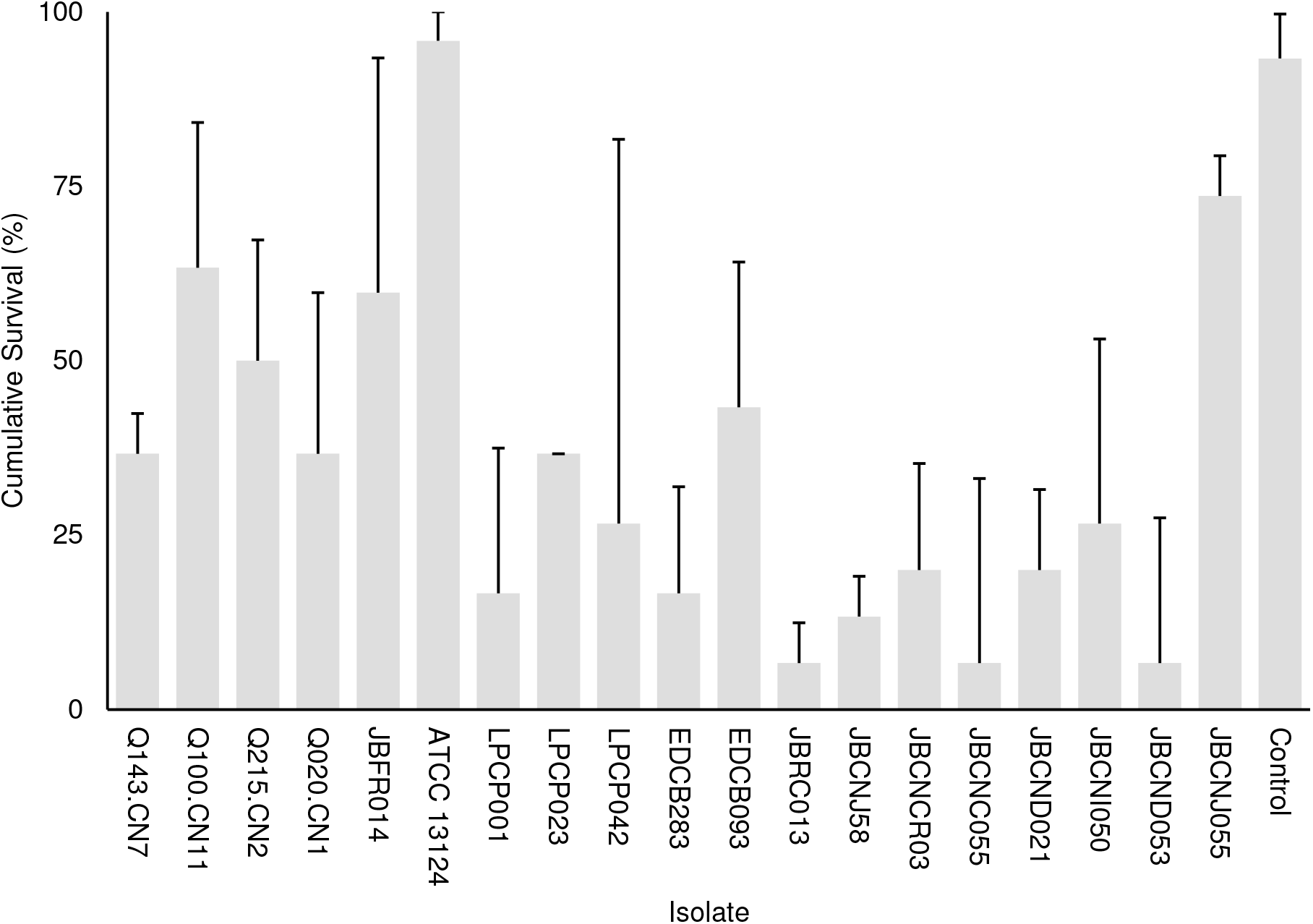
Average survival percentage after 72 hours incubation for *G. mellonella* infected with 10μL of 10^7^ CFU/mL of respective isolates. Isolates derived from human sources show a higher degree of pathogenicity to those from human sources. Error bars indicate standard deviation.

Parental injection of 10^5^ CFU JBCNJ055 *C. perfringens* resulted in disease of the larvae. Larvae which succumb to the infection exhibit nodulation, blackening of the cuticle and eventually death. Figure 3 shows the rate of development of infection appears dependent on inoculum size as melanisation score increases rapidly with increasing inoculum doses of JBCNJ055. Kaplan-Meier survival distributions for each bacterial inoculum of JBCNJ055 were significant when compared using the log-rank (Mantel-Cox) test (pooled over strata) (P<0.001). Survival probability appears dependent on the number of organisms injected (Figure 4). An inoculum size of 10^5^ CFU/larvae was required for 73.3 ± 6.66% survival and induced melanisation (>3) in >80% of the population. 10^4^ CFU/larvae produced 79.2 ± 7.2% survival however <30% of the population exhibit melanisation scores higher than 2. Further dilutions (10^3^; 10^2^; 10) produced >90% survival. Injection with 10^5^ CFU/larvae of JBFR014 did incite melanisation in ~40% of the population however survival remained at >90%. Injection with lower dilutions (10^3^; 10^2^; 10) resulted in no visible blackening of the larvae. Interestingly, injection with 10ul of 10^7^ CFU/mL ATCC^®^ 13124™ *C. perfringens* did not cause visible disease to the larvae within the 72 hour window investigated. Melanisation did occur at the site of injection however no further immune response was visually observed. Survival remained above 95% at each inoculum group investigated.

**Figure 3:**
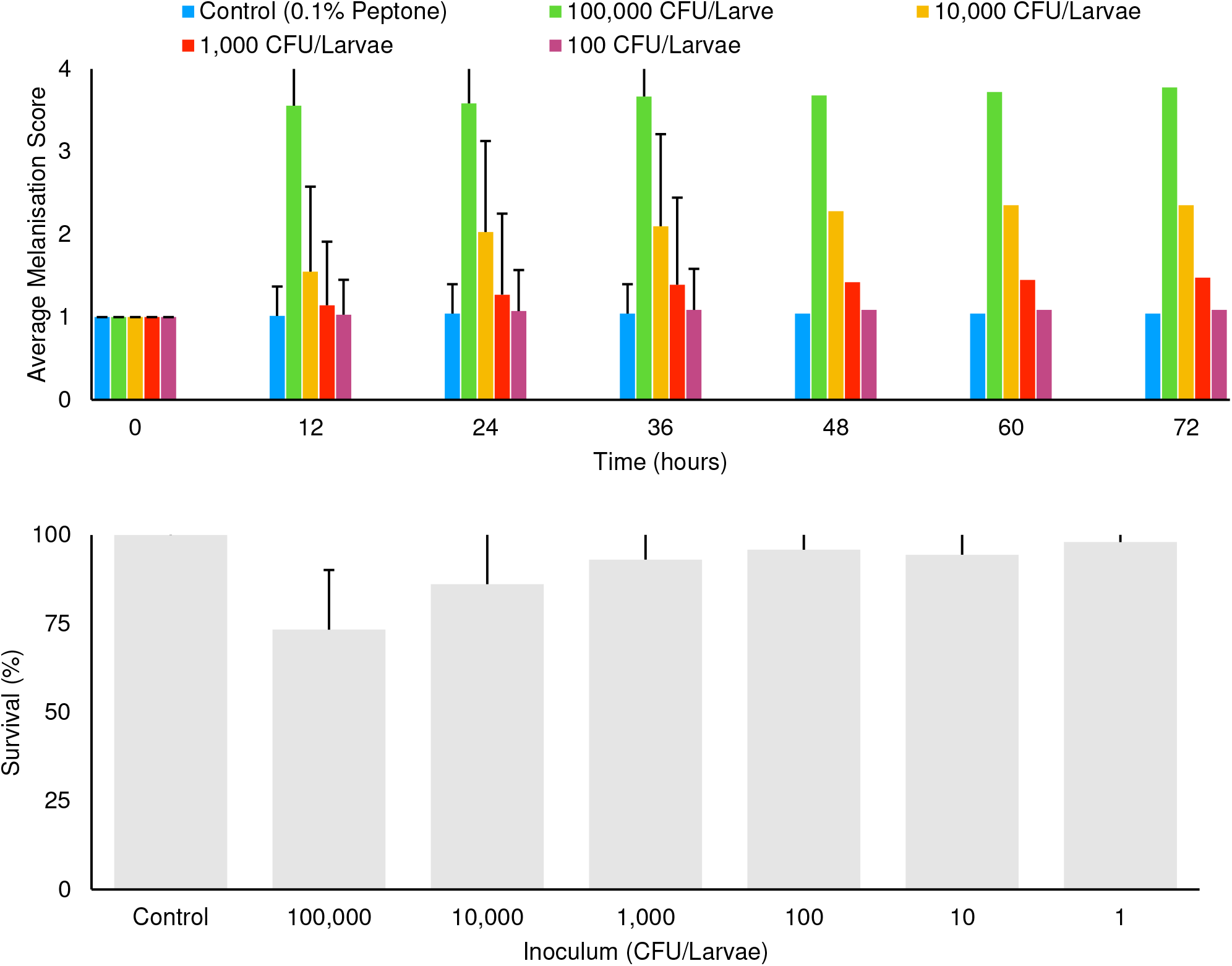
Mean melanisation score of pooled data, 72 hours post inoculation of 10^7 -^ 10^3^ CFU/mL of *Clostridium perfringens* JBCNJ055. Scoring system is as follows: 1= healthy, 2= <50% melanisation, 3= >50% melanisation, 4 = deceased. Data shown is pooled from three distinct repeats and error bars represent standard deviation. Injection with 10^7^ JBCNJ055 was required to cause potent melanisation resulting in scored higher than 3. Survival percentages of *G. mellonella* after inoculation with 10^7 -^ 10^3^ CFU/mL of *Clostridium perfringens* JBCNJ055. An injection with 10^7^ of JBCNJ055 was also required to result in a substantial fall in larval survival. Error bars show standard deviation.

**Figure 4:**
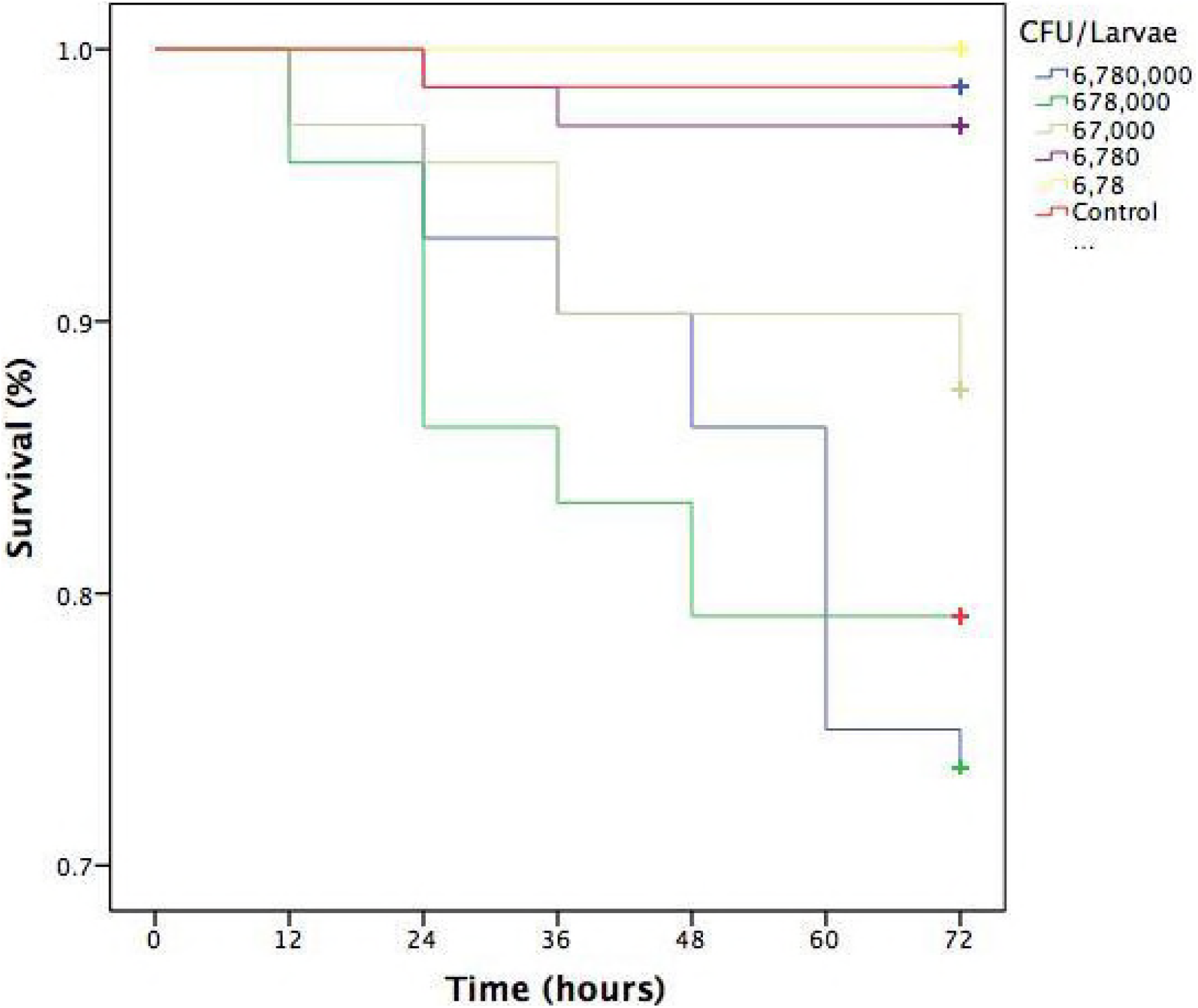
Kaplan-Meier survival distributions for dose dependant challenge of JBCNJ055. Three repeats of the same experiment were pooled. Results were collated as % survival. Strikethrough indicates censored data (unaffected larvae). All levels of infection data are significant (P<0.001) indicating larval survival is based upon the amount of bacteria injected.

### Phenoloxidase

The results of the phenoloxidase assays are shown in Figure 5 and shows differences between the control and inoculated groups but not at a significant difference level. Infected groups demonstrated a melanisation score of 3 ± 0.8 at 48 hours post inoculation compared to no melanisation in 0.1% peptone controls.

**Figure 5:**
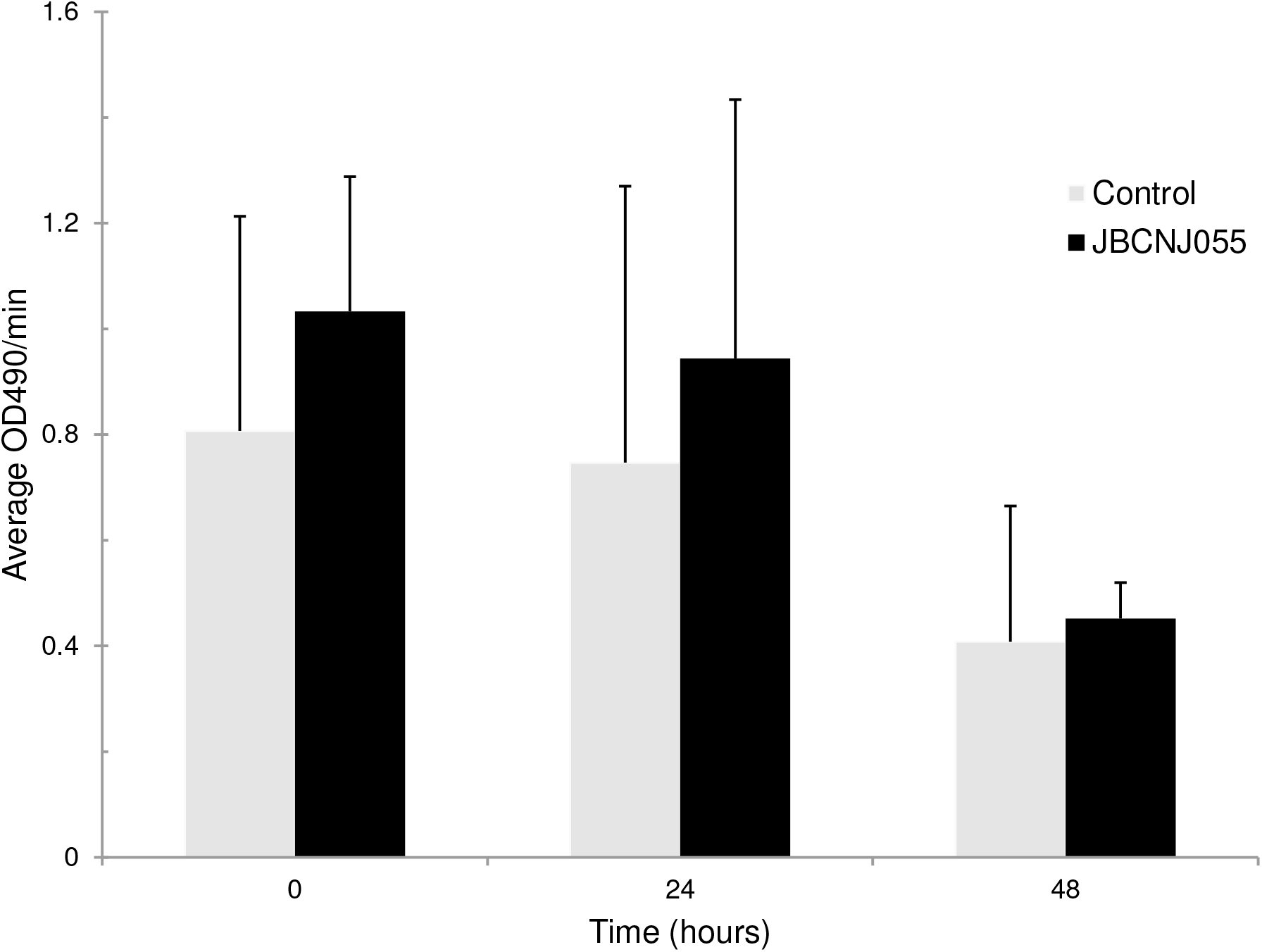
Phenol oxidase enzyme activity assay adapted from Eleftherianos *et al*. (2008). Haemolymph from larvae infected with 10μL of 10^7^ JBCNJ055, was assayed for phenol oxidase activity and compared to controls. Error bars show standard deviation. Slight differences can been seen between infected larvae and controls although not significant. Indicates the activation of the phenol oxidase pathway but no increase in enzyme expression.

### Histopathology of Galleria samples

Figure 6 shows a distinct response to a pathogen compared to controls demonstrating aggregation and nodulation of haemocytes. There is clear loss of structure and tissue damage in samples which had been injected with 10^5^ CFU/larvae of JBCNJ055, with an abundance of melanin demonstrated compared with negative controls. Gram-positive were observed associated with tissue remnants.

**Figure 6:**
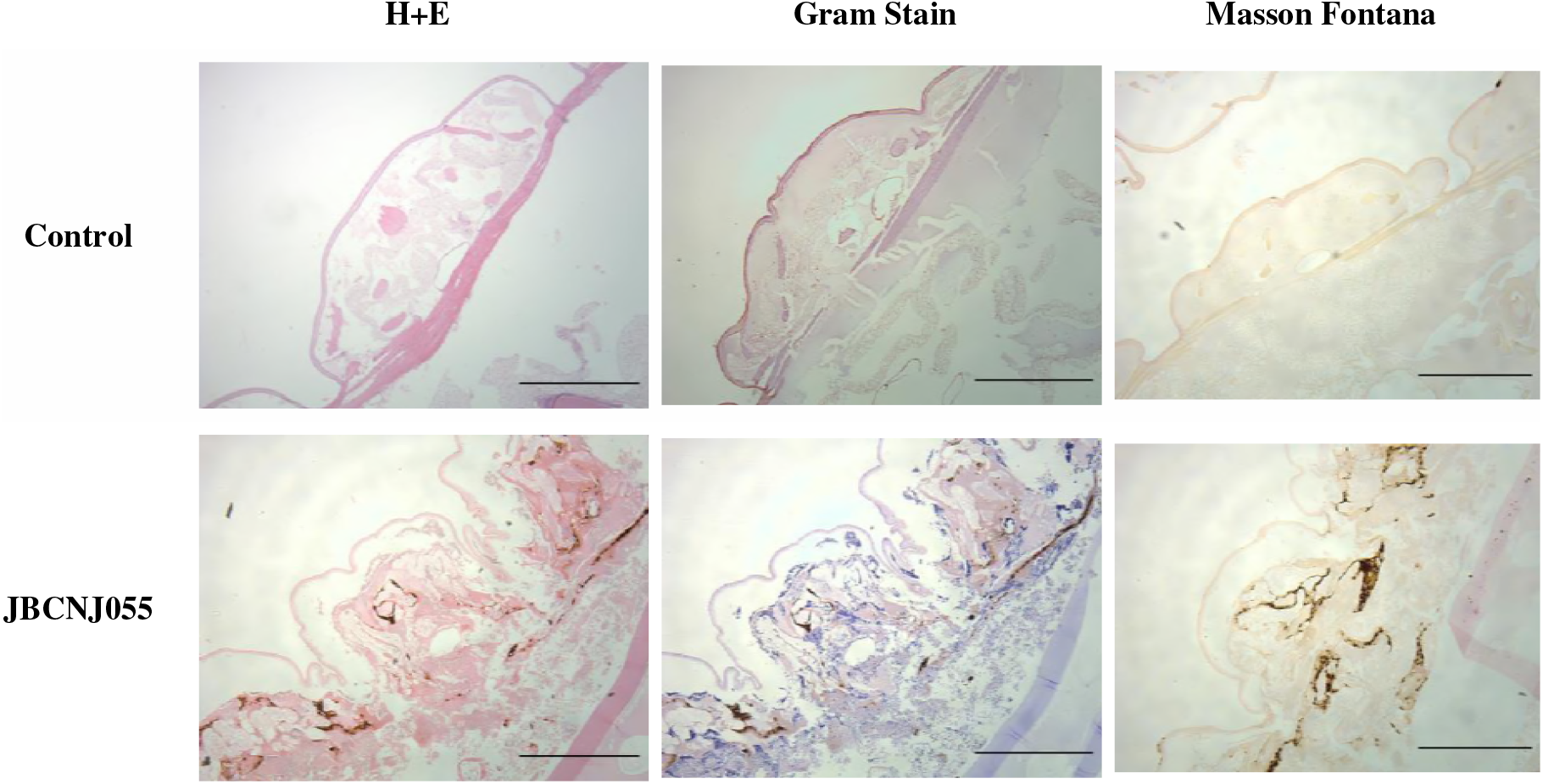
**(x40)** Comparison of *G. mellonella* histopathology of control and infected groups after 72 hours incubation. Sections were stained with H+E, Gram stain and Masson Fontana (highlight melanin deposition). Infected groups show distinct loss of tissue structure, systemic proliferation of Gram-positive bacteria and a large production of melanin within peripheral tissues indicative of systemic pathogenicity. Scale bars represent 1mm.

### Antibiotic therapy

No melanisation or death of larvae was observed with any antibiotics in the absence of infection prior to therapy trials (data not shown). Table 3 shows penicillin G to be the most effective agent *in vitro* against JBCNJ055 (MIC: 0.18μg/mL) however *in vivo* larval survival was reduced by 7.5% compared with controls. Interestingly, Figure 7 shows that the penicillin G treatment resulted in a rapid onset of melanisation compared with controls. We have observed a potent systemic melanisation in >80% of the larvae in under 60 minutes after the treatment dose was administered. Injection of penicillin G alone was not sufficient to cause this potent reaction. Both tetracycline and bacitracin were effective against *C. perfringens* JBCNJ055 *in vitro* (MIC: 1.56μg/mL). *In vivo*, tetracycline increased larval survival by 7.5% whereas bacitracin greatly increased larval survival by a remarkable 28.3% making it the most effective agent *in vivo* (Table 3). Unsurprisingly, neomycin was the least effective *in vitro* (MIC: 60μg/mL) alongside a reduction in larval survival *in vivo* of 11.3%.

**Figure 7:**
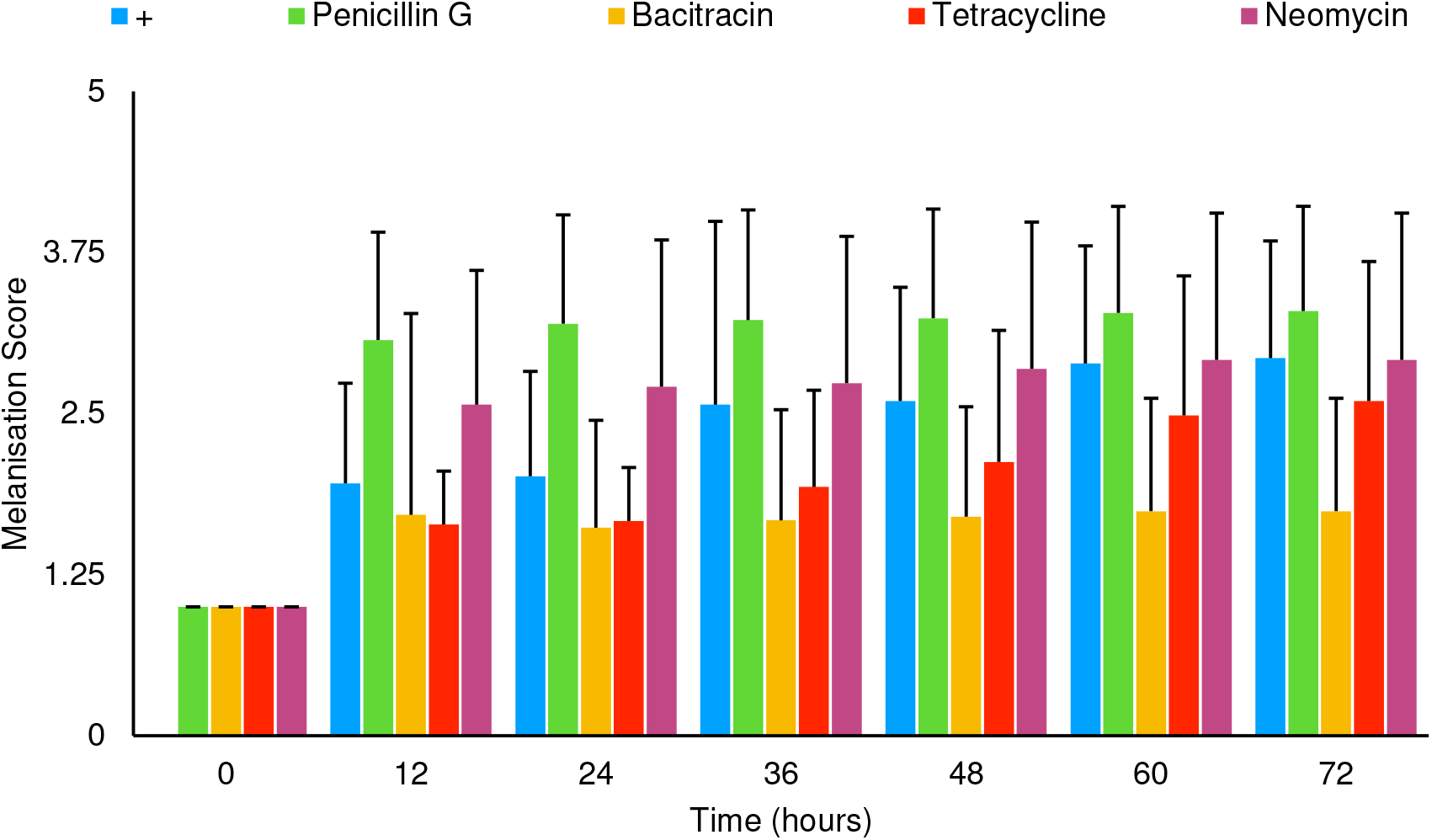
Average melanisation Score for larvae infected with 10μL of 10^7^ JBCNJ055 with a secondary injection of either penicillin G (2mg/kg), bacitracin (64mg/kg), tetracycline (64mg/kg), or neomycin (2400mg/kg). healthy larvae with no signs of infection were scored 1. Larvae showing nodulation and lateral line melanisation scored 2. Larvae exhibiting systemic melanisation scored 3. Deceased, fully pigmented larvae scored 4. There is a potent increase in melanisation score when penicillin G is used in response to the infection. Likely due to oxidative stress in the midgut predisposing larvae to disease.

## DISCUSSION

*G. mellonella* is becoming a well established *in vivo* model for virulence studies of a number of important microbes and is gaining momentum in the literature as more pathogens are investigated. The advantages of an insect model over mammalian studies in terms of ethics, speed and research cost are becoming apparent with increasing data (Jander *et al.*, 2000). The time lapse features developed here allows for reduced labour inputs although it is not possible to physically manipulate the larvae, a common practice in assessing larval mortality (Peleg *et al.*, 2009; Harding *et al.*, 2013; Beeton *et al.*, 2015). This has been overcome in the present study by classifying dead insects by a melanisation score of 4 (Table 4) coupled with no movement for 20 minutes (2 frames). Time-lapse methodologies however increase precision by reducing intermittent 12 hour observations to 10 minute increments. The increase in precision is of benefit when studying pathogens with rapid disease progression. Inherent bias in the manual scoring of the video should be noted but this can be reduced by our current development of an automated melanisation recognition system utilising the time-lapse feature which will allow for a more robust and standardised scoring system and reduced processing time.

### *Galleria mellonella* and *Clostridium perfringens*

*C. perfringens* is a ubiquitous organism, with a potential arsenal of toxins and hydrolytic enzymes associated with pathogenicity (Petit *et al.*, 1999). The anaerobic pathogen is economically devastating to the global food industry and is estimated to cost the poultry sector alone over $6 billion (USD) per year (Wade and Keyburn, 2015). We have shown for the first time *C. perfringens* is pathogenic towards *G. mellonella* larvae although the degree of pathogenicity is distinct between isolates. The differences in pathogenicity of isolates may be due to differing toxin expression or unaccounted environmental factors which are thought to play a major role in mammalian CPAD (Borda-Molina *et al.*, 2018). Equivalent inocula of *C.perfringens* were generally required to produce larval death which was consistent with other bacteria that have been studied such as *Campylobacter jejuni* (10^6^ CFU/larvae *Yersinia pseudotuberculosis* (10^6^ CFU/larvae) and *Clostridium difficile* (10^5^ CFU/per larvae) (Senior *et al.*, 2011; Champion *et al.*, 2009; Nale *et al.*, 2015 respectively).

CPADs are complex and multifactorial processes which are still poorly understood (Silva and Loboto, 2015). Although advancements have been made, the use of the larval model offers a viable alternative approach for high throughput with radical and novel therapeutic agents. Similar pathogenic studies have included immunological aspects of infection such as haemocyte viability studies (Bergin *et al.*, 2003) and a variety of histopathological approaches (Harding *et al.*, 2013) to further the understanding of pathogenic processes. In the present study, *C.perfringens* does activate the phenoloxidase pathway although our enzyme activity data suggests no upregulation of the enzyme in response as seen with other pathogens (Harding *et al.*, 2012). Broader studies are needed to draw conclusions about the mechanism of pathogenicity with the larvae and to identify what important environmental confounding elements exist.

Treatment of the infection with antimicrobials appears to broadly correlate with the *in vitro* data obtained. Penicillin G, surprisingly induces a robust and potent reaction as a response to infection, which was not seen when the antibiotic was administered alone. Previous studies have highlighted that penicillin G induces high levels of oxidative stress in the midgut of *G. mellonella* (Büyükgüzel and Kalender, 2007) which may predispose the larvae to disease. Neomycin was the least effective agent and the reduction in larval survival is likely due to disruption of native microbiota allowing for proliferation of opportunistic pathogens with little or no effect on invading *C. perfringens*. Tetracycline and bacitracin are both equally effective *in vitro* against JBCNJ055 but the efficacy of bacitracin *in vivo* is likely due to the more bactericidal nature of bacitracin against Gram-positive organisms in comparison to the broad spectrum but bacteriostatic tetracycline.

*G. mellonella* is becoming a powerful alternative model for the investigation of bacterial pathogenicity and we have shown for the first time its use in the investigation of *C. perfringens* infection. The mode of pathogenicity of *C. perfringens* in *G. mellonella* remains elusive and further investigation is required to determine the factors surrounding the progression of the disease. We have also devised a methodological advancement which may increase the speed of research further. We consider that the larvae model is unlikely to replace the mammalian model in physiological studies of disease or pathogenesis but will be a useful development to characterise pathogen virulence in important strains and perhaps allow pre-screening of antimicrobials against pathogens before embarking on expensive and demanding mammalian clinical trials.

## Acknowledgements

Our thanks to Emily Brandreth, Luke Randall and Danielle Bramley for their technical contributions.

